# Effect of developmental and adult diet composition on reproductive aging in *Drosophila melanogaster*

**DOI:** 10.1101/2024.05.14.589018

**Authors:** B G Ruchitha, Devashish Kumar, Mohankumar Chandrakanth, Itibaw Farooq, Nishant Kumar, Chand Sura, S. Chetan, Sudipta Tung

## Abstract

Diet significantly affects reproductive outcomes across species, yet the precise effects of macronutrient compositions beyond caloric intake on reproductive aging are understudied. Existing literature presents conflicting views on the fertility impacts of nutrient-rich versus nutrient-poor developmental diets, underscoring a notable research gap. This study addresses these gaps by examining effects of isocaloric diets with varied protein-to-carbohydrate ratios during both developmental and adult stages on reproductive aging of a large, outbred Drosophila melanogaster population (n = ∼2100). Our results clearly demonstrate an age-dependent dietary impact on reproductive output, initially dominated by the developmental diet, then by a combination of developmental and adult diets in early to mid-life, and ultimately by the adult diet in later life. Importantly, we found that the effects of developmental and adult diets on reproductive output are independent, with no significant interaction. Further investigations into the mechanisms revealed that the effect of developmental diet on fecundity is regulated via ovarioles formation and vitellogenesis; while, the effect of adult diet on fecundity is mostly regulated only via vitellogenesis. These insights resolve disputes in the literature about dietary impacts on fertility and offer valuable perspectives for optimizing fertility strategies in improving public health and conservation efforts in this changing world.

**Highlights:**

1. Effect of developmental and adult diet composition on reproduction is age-dependent
2. Developmental diet affects early-life; adult diet late-life; and both affect mid-life
3. But the effect of developmental and adult diets do not interact with each other
4. Developmental diet regulates reproduction via ovarioles formation and vitellogenesis
5. Whereas, adult diet regulates reproduction via differential vitellogenesis across age

## INTRODUCTION

Environmental conditions during development influence the expression of traits in adult life. Particularly, the diet consumed during development impacts the individual’s health throughout their life (Grafen, 1988; Hales and Barker, 1992; Madsen and Shine, 2000; Gluckman and Hanson, 2004; Van De Pol *et al*., 2006; Millon *et al*., 2011; Black *et al*., 2013; Wong and Kölliker, 2014; May *et al*., 2015; Pigeon *et al*., 2019; Duxbury and Chapman, 2020; Spagopoulou *et al*., 2020). Thus, several attempts have been made to understand the effect of developmental diet on adult traits. In 1988, based on inferences drawn from several fertility studies where early food availability was documented (Sadler, 1969), Grafen proposed the silver spoon hypothesis. The silver spoon hypothesis posits that organisms reared on a better quality developmental diet have better fitness traits as adults (Grafen, 1988). Soon after, in 1992, Hales and Barker proposed the thrifty phenotype hypothesis to explain the occurrence of type 2 diabetes in humans as a mismatch of diets during developmental and adult stages of life. The thrifty phenotype hypothesis states that individuals who experience poor nutrition during developmental stages, program their metabolism with the expectation of poor nutrition in adult life. And so, if the quality of nutrition improves during adult life, the individuals’ adaptation to poor nutrition now becomes maladaptive and they suffer from metabolic disorders like type 2 diabetes (Hales and Barker, 1992). Following this, in 2004, Gluckman and Hanson proposed a more generalized version of the thrifty phenotype hypothesis called the predictive adaptive responses (PAR). PAR states that irrespective of the shift in developmental diet to adult diet being poor to rich or rich to poor, the individuals will suffer a fitness cost in adult life due to maladaptation to the change in adult diet as compared to their developmental diet (Gluckman and Hanson, 2004). Since the inception of these hypotheses, various studies have been published in support of the silver spoon as well as the thrifty phenotype and the PAR hypotheses (Madsen and Shine, 2000; Van De Pol *et al*., 2006; Millon *et al*., 2011; Wong and Kölliker, 2014; Pigeon *et al*., 2019; Duxbury and Chapman, 2020). However, particularly in recent years, there have been reports of instances in which the observed effects of developmental environment conditions on adult traits cannot be explained by the silver spoon or the thrifty phenotype/PAR hypotheses (May *et al*., 2015; Spagopoulou *et al*., 2020).

This has led to several studies that attempt to explain the above-stated disagreement in the effect of developmental diet on adult traits by highlighting the context dependencies of the above hypotheses. One such study (May *et al*., 2015) in *Drosophila melanogaster* empirically shows that the effect of developmental diet on adult traits varies depending on adult diet and mating status. They demonstrate that when the adult diet is supplemented with yeast (source of protein) and they have continuous access to mates, populations reared on a rich developmental diet have decreased reproductive output as compared to populations reared on a poor developmental diet i.e. the silver spoon and the thrifty phenotype/PAR hypothesis cannot explain the effect of developmental diet on adult reproductive output when the adult reproductive potential is not constricted by unavailability of mates or dietary proteins (May *et al*., 2015). Further, in 2020, Duxbury and Chapman find that the effect of developmental diet on reproductive output is seen only in females, and not in males. Even within females, they see that the silver spoon and thrifty phenotype/PAR hypotheses hold true when adult diet is high in protein; and only the silver spoon hypothesis holds true when adult diet is low in protein (Duxbury and Chapman, 2020). Adding to this body of knowledge, in this study, we have attempted to further explore the nuances in how developmental and adult diets affect reproductive output in adult life.

Reproductive ability varies with age (May *et al*., 2015; Duxbury and Chapman, 2020). Thus the effect of developmental and adult diets on reproductive ability of an organism would also vary with age. Additionally, reproduction is affected by various aspects of the diet like calorific value, macronutrient composition, etc. (Lee *et al*., 2008). However, studies that look at the effect of developmental and adult diets on reproductive output change both calorific value and macronutrient composition of the diet simultaneously (May *et al*., 2015; Duxbury and Chapman, 2020). Therefore, one cannot distinguish between the effects of the two. In our study, we used isocaloric dietary combinations of varying macronutrient compositions provided at developmental and adult life stages in *Drosophila melanogaster* populations to investigate how reproductive ability varies with age in response to dietary macronutrient changes. Using isocaloric diets removes the confounding effect of the calorific value to the diets that vary in macronutrient composition. We only vary the two major macronutrients – proteins and carbohydrates, and use a full-factorial combination of the diets applied to developmental vs. adult life stages. We use this system to examine the effects of the developmental and adult dietary combinations on the number of viable progeny produced in early, mid and late adult life. We see that the effects of developmental diet on reproductive ability is independent of adult diet. We observe the silver spoon theory is true in early-life (i.e. high protein developmental diet results in more viable progeny, irrespective of the adult diet); while, neither silver spoon nor the thrifty phenotype/PAR hypotheses are applicable in mid-life (i.e. low protein developmental diet results in more viable progeny, irrespective of the adult diet); and, the developmental diet does not affect the number of progeny produced in late-life.

To further understand the underlying physiological basis of these differences, we examine how these dietary combinations affect the process of egg production since ovary formation and oogenesis regulation are known to be affected by inputs from nutrient-sensing pathways (Bownes, 1989; Green and Extavour, 2014). Developmental diet is known to affect reproduction in adult life via formation of ovarioles during larval stage (Hodin and Riddiford, 2000; Tu and Tatar, 2003). Our results show that development of ovarioles alone does not explain the age-dependent effect of developmental diet on reproductive output. Although recent studies, by examining gene expression, have shown that the effect of developmental diet on reproduction is mainly restricted to early adult life (May *et al*., 2021; Collins *et al*., 2023), the physiological basis of this early-life regulation of reproduction via developmental diet is unknown and thus the link from changes in gene expression to variation in offspring production via changes in diet is not well understood. Furthermore, there exist very few studies that look at how developmental and adult diets interact to shape adult traits like fecundity (May *et al*., 2015; Macartney *et al*., 2017, 2018; Duxbury and Chapman, 2020).

Our results, by examining the anatomical and physiological basis of regulation of the process of egg production via developmental and adult diets in an age-dependent manner and mapping it back to the number of progeny produced by populations reared in these dietary conditions, advance our understanding of dietary effects on phenotypic plasticity of reproduction and reproductive aging.

Overall, the results from our study highlight the age-dependent nature of the effect of developmental and adult diets on adult reproductive output. Additionally, by examining the underlying physiological processes involved in reproduction in *D. melanogaster* females, we also show that although the number of progeny produced is dependent on the effect of both developmental and adult diets across age, the various physiological processes involved in egg production are not necessarily dependent on both the diets. Specifically, the number of ovarioles is dependent only on developmental diet; while, ovary size and the proportion of egg chambers that undergo vitellogenesis are dependent on both developmental and adult diets, and vary with age. These results address some of the key conflicts that exist in literature about the effect of developmental diets on adult reproductive output and highlight the importance of measuring adult traits in an age-specific manner.

## MATERIALS AND METHODS

### 2.1. Experimental population

We used a large (n = ∼2100) outbred population of *Drosophila melanogaster* reared on a corn-sugar-yeast based diet of protein to carbohydrate ratio of 0.4 for at least 35 generations. This baseline population was derived from offsprings produced by random mating of flies from four MB populations (Mudunuri *et al*., 2024). The baseline population was reared at a density of ∼300 eggs per 50mL of food. Adults eclosing from seven such replicates were pooled together into a plexi-glass cage (25□cm□×□20□cm□×□15□cm) where a egg-laying surface would be provided for 12 to 16 hours. In this manner, the baseline population was maintained in a 14-day generation cycle in constant light at 25°C. For the experiments, progeny of this baseline population were subjected to four experimental isocaloric diet regimes with varying protein to carbohydrate ratios across developmental and adult stages.

Experimental flies were reared on an uncrowded density of ∼150 eggs per 50 mL of developmental diet. All assays were performed on flies from these experimental regimes by sampling the population cross-sectionally.

### 2.2. Experimental diet

The ancestral diet is a corn-sugar-yeast media with a protein to carbohydrate ratio of 0.4. For a high protein-to-carbohydrate diet, we used a protein to carbohydrate ratio of 0.7; and for a low protein-to-carbohydrate diet, we used protein to carbohydrate ratio of 0.25 (Mudunuri *et al*., 2024). All the diets used are isocaloric in nature. Detailed information on the diets can be found in Mudunuri *et al*., 2024.

The ancestral baseline population was raised on a diet of protein to carbohydrate ratio of 0.4 in both developmental and adult stages. The experimental flies were subjected to four experimental isocaloric diet regimes with varying protein to carbohydrate ratios across developmental and adult stages – HH, HL, LH, LL. Here, H represents high protein-to-carbohydrate diet (henceforth, high protein diet) and L represents low protein-to-carbohydrate diet (henceforth, low protein diet), and the two-letter code indicates the protein level of developmental and adult diet respectively.

### 2.3. Assays

#### 2.3.1. Viable progeny count

To estimate reproductive output, two males and two females were aspirated from the adults reared in plexiglass cages into a glass vial containing ∼5 mL of adult food on the day of the experiment. They were then allowed to oviposit for 24 hours. There were 30 such vial replicates per treatment. After the egg-laying window, replicates with dead females were discarded and the remaining vials were then incubated for 13 days to give sufficient time for the eggs to develop into adults. The enclosed adults were counted to quantify the number of viable progenies.

#### 2.3.2. Ovary size measurement

To measure ovary size, ovaries of 30 females from each treatment were dissected in 1X Phosphate-buffered saline (PBS; SRL, #78529). The images of the dissected ovaries were captured immediately under a stereomicroscope (Zeiss, Stemi 305) to determine the size of the ovaries. To quantify the size of the ovary, Region of Interest (ROI) was drawn around the ovary and area of the ROI was measured in ImageJ2.

#### 2.3.3. Ovariole count

To quantify the number of ovarioles, ovaries of 30 females per treatment were dissected in 1X PBS. The dissected ovaries were placed in 4% Paraformaldehyde (PFA; Sigma-Aldrich, #P6148-500G) solution for 15 minutes to reduce the chances of breakage of ovarioles during further dissection. The ovaries were then shifted back to 1X PBS, and the ovarioles of each ovary were separated and counted under light microscope to quantify the number of ovarioles per female.

#### 2.3.4. Egg chambers vitellogenesis quantification

To estimate the proportion of egg chambers that undergo vitellogenesis, ovaries of 20 females per treatment were dissected in 1X PBS. The dissected ovaries were placed in 4% PFA solution for 15 minutes. The ovaries were then shifted back to 1X PBS, and the ovarioles separated. Egg chambers in pre-vitellogenesis and those that undergo vitellogenesis were counted under light microscope. Oocyte being larger than the largest nurse cell was taken as the marker for vitellogenesis (Bakken, 1973). So, all egg chambers with oocytes smaller than the largest nurse cell were binned as pre-vitellogenesis egg chambers; whereas, all egg chambers with oocytes larger than the largest nurse cell were binned as egg chambers that undergo vitellogenesis.

Due to the logistical constraints posed by large sample sizes for each assay, the flies used for each of the above-mentioned assays were collected through separate egg collection rounds and maintained on specific treatment diets.

### 2.4. Experiments

#### 2.4.1. To investigate the effect of developmental and adult diets on age-dependent reproductive output

To understand how developmental and adult nutrition interact across age to influence reproductive output, we counted the number of viable progeny produced by flies reared in HH, HL, LH and LL diets in very-early, early, mid and late ages of adult life i.e. on days 10, 12, 22 and 32 from the day of egg collection. On the day of the experiment, two male-female pairs from individuals reared in plexiglass cages under respective treatment dietary regimes were aspirated into vials containing fresh adult food. They were allowed to oviposit for 24 hours, eggs were reared under standard environmental conditions, and the resulting viable progeny were then counted. We set up 30 such vial replicates for each treatment. In total, we counted viable progeny produced by 960 male-female pairs for this experiment – 30 vials × 2 male-female pairs × 4 dietary regimes × 4 age-points.

The last age point i.e. day 32 was chosen as a day before the start of demographic senescence in the population (Mudunuri *et al*., 2024) to avoid survivorship bias in the data. Day 10 represents the day post-eclosion peak under all diet regimes. Day 12 represents the early-life fecundity peak of the populations (preliminary observation). Day 22 was chosen as the mid-point between Day 12 and Day 32. In this article, we refer to age 10, 12, 22 and 32 days post-egg collection as very-early, early, mid and late-life respectively and use them interchangeably.

#### 2.4.2. To test the contribution of high early-life reproduction on decrease in mid-life reproductive output in populations that consumed high protein adult diet

Based on the result that in case of varying developmental diet and high protein adult diet, there exists a tradeoff in reproductive output between early-life and mid-life (Figure 1A; Table S2; Table S4), we hypothesized that high early-life reproduction causes the decrease in mid-life reproductive output in populations that were reared in HH dietary regime as opposed to LH dietary regime. To test this hypothesis, we reared the flies from the high protein adult diets (HH and LH) as virgins for 20 days to remove the effect of early-life reproduction, and then allowed them to mate and reproduce in mid-life for a day. We then counted and analyzed the number of viable progeny produced.

**Figure 1.**
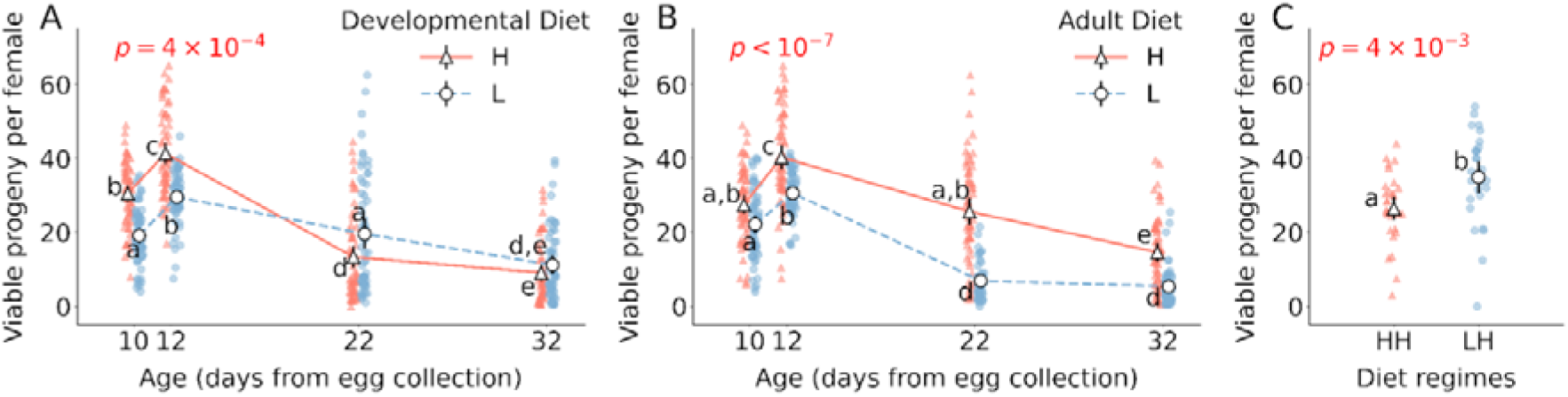
Effect of developmental and adult diet on reproductive output. (A) The effect of developmental diet across age on the number of viable progeny produced. (B) The effect of adult diet across age on the number of viable progeny produced. (C) The number of viable progeny produced by populations reared as virgins on HH and LH dietary regimes in early-life and allowed to mate and reproduce in mid-life. The triangle marker with solid line represents high protein diet H; the circle marker with dotted line represents low protein diet L. The black solid vertical lines indicate 95% confidence interval for each treatment. Some error bars are not visible because of their small size. The p-value in (A) is of developmental diet × age interaction effect. The p-value in (B) is of adult diet × age interaction effect. Different lower-case letters denote statistically significant differences (*p* < 0.05).

If mid-life reproductive output does trade-off with early-life reproductive output, then the difference between HH and LH in reproductive output in mid-life should no longer exist when investment into reproductive output in early-life has been minimized.

#### 2.4.3. Understanding the physiological basis of the differential impact of development and adult diets on age-dependent reproductive output

To understand how developmental and adult nutrition interact across age to influence reproductive ability, we measured the size of ovaries, counted the number of ovarioles, and quantified the efficiency of vitellogenesis of egg chambers in flies reared in HH, HL, LH and LL diets in very-early, early, mid and late phases of adult life i.e. on days 10, 12, 22 and 32 from the day of egg collection. In total, for these experiments, we dissected 1,280 females for the three assays – 30 females × 4 dietary regimes × 4 age-points for measuring ovary size, 30 females × 4 dietary regimes × 4 age-points for quantifying the number of ovarioles, and 20 females × 4 dietary regimes × 4 age-points for quantifying egg chamber stages.

### 2.5. Statistical analysis

#### 2.5.1. The effect of developmental and adult diets on age-dependent reproductive output

To analyse viable progeny count data, we used generalised linear models (GLMs) with negative binomial error structure and log link function to investigate the relationship between the three fixed factors (with respective levels) – developmental diet (H and L), adult diet (H and L), age (10, 12, 22, 32), and the response variable – viable progeny count. When the interaction effect was significant, post-hoc was performed using Tukey’s HSD test.

To understand how the different combinations of developmental and adult diets compare with one another, we calculated effect size using Cohen’s d (Cohen, 1988) for all treatment comparisons within each age point. Cohen’s d value less than 0.2 was considered non-significant, between 0.2 and 0.5 was considered as small effect size, between 0.5 and 0.8 was considered as medium effect size, more than 0.8 was considered as large effect size.

#### 2.5.2. The contribution of high early-life reproduction on decrease in mid-life reproductive output in populations that consumed high protein adult diet To test whether decrease in mid-life reproductive output is due to greater early-life reproduction in populations that consumed a high protein adult diet, we counted the number of viable progeny in flies reared as virgins in early-life in HH and LH dietary regimes, and allowed them to mate and reproduce only in mid-life. We used generalised linear models (GLMs) with negative binomial error structure and log link function to investigate the relationship between the fixed factor (with respective levels) – treatments (HH and LH), and the response variable – viable progeny count.

#### 2.5.3. Physiological basis of differential reproductive output

##### 2.5.3.1. Ovary size

To analyse ovary size data, we used generalised linear models (GLMs) with Gamma error structure and identity link function to investigate the relationship between the three fixed factors (with respective levels) – developmental diet (H and L), adult diet (H and L), age (10, 12, 22, 32), and the response variable – ovary size. When the interaction effect was significant, post-hoc was performed using Tukey’s HSD test.

##### 2.5.3.2. Number of ovarioles

To analyse the data of the number of ovarioles, we used generalised linear models (GLMs) with negative binomial error structure and log link function to investigate the relationship between the three fixed factors (with respective levels) – developmental diet (H and L), adult diet (H and L), age (10, 12, 22, 32), and the response variable – number of ovarioles.

##### 2.5.3.3. Vitellogenesis of egg chambers

To analyse data of the total number of egg chambers, we used generalised linear models (GLMs) with negative binomial error structure and log link function to investigate the relationship between the three fixed factors (with respective levels) – developmental diet (H and L), adult diet (H and L), age (10, 12, 22, 32), and the response variable – number of egg chambers. Post-hoc was performed using Tukey’s HSD test.

To understand how developmental and adult nutrition interact across age to affect vitellogenesis of egg chambers, we calculated the proportion of egg chambers that undergo vitellogenesis (i.e. number of egg chambers post vitellogenesis/ total number of egg chambers) in flies reared in HH, HL, LH and LL diets on days 10, 12, 22 and 32 from the day of egg collection. We used generalised linear models (GLMs) with gamma error structure and log link function to investigate the relationship between the three fixed factors (with respective levels) – developmental diet (H and L), adult diet (H and L), age (10, 12, 22, 32), and the response variable – proportion of egg chambers that undergo vitellogenesis. When the interaction effect was significant, post-hoc was performed using Tukey’s HSD test.

All data was analyzed using R 4.3.2; and all plots were created in Python 3.11 using Spyder Version 5.

## RESULTS

### 3.1. The effect of developmental and adult diets on reproductive output is dependent on age

Effect of developmental diet on reproductive output, measured as the number of viable progeny produced by a male-female pair, varies significantly with age (developmental diet × age interaction F_3,455_ = 6.1, *p* = 4×10^−4^; Figure 1A; Table S1). In very-early and early-life, populations reared on a high protein developmental diet produce more progeny as compared to populations reared in low protein developmental diet (for 10th day Tukey’s HSD *p* = 4.4 ×10^−7^, for 12th day Tukey’s HSD *p* = 2×10^−3^; Figure 1A; Table S2). However, interestingly, in mid-life, populations reared on a low protein developmental diet produce more progeny as compared to populations reared on a high protein developmental diet (Tukey’s HSD *p* = 0.01; Figure 1A; Table S2). In late-life, developmental diet has no effect on the number of progeny produced (Tukey’s HSD *p* = 0.64; Figure 1A; Table S2).

Based on the above result, there appears to exist a tradeoff between early-life and mid-life reproductive output. Populations that are reared on a high protein developmental diet have higher reproductive output in early-life and have lower reproductive output in mid-life in comparison to populations reared on low protein developmental diet (Figure 1A; Table S2).

Similarly, the effect of adult diet on the number of viable progeny produced varies significantly with age (adult diet × age interaction F_3,455_ = 21.7, *p* < 10^−7^; Figure 1B; Table S1). Populations reared on high protein adult diet produce more progeny than populations reared on low protein adult diet at all measured age points, except in very-early-life i.e. 10th day from egg collection (for 10th day Tukey’s HSD *p* = 0.11, for 12th day Tukey’s HSD *p* = 0.04, for 22nd day Tukey’s HSD *p* < 10^−7^, for 32nd day Tukey’s HSD *p* < 10^−7^; Figure 1B; Table S3).

By looking at the effect size of pairwise comparisons of the treatments within each age point, we see that the tradeoff between early-life and mid-life reproductive output when developmental diet is changed is driven by populations reared in high protein adult diet rather than by populations reared in low protein adult diet (i.e. although in early-life HH > LH and HL > LL with large effect size, in mid-life LH > HH with medium effect size and LL > HL with small effect size; Table S4).

Furthermore, the effect of developmental diet on viable progeny numbers is independent of adult diet and vice versa (developmental diet × adult diet interaction F_1,455_ = 0.3, *p* = 0.6; Table S1). The three-way interaction of developmental diet, adult diet and age is also not significant (developmental diet × adult diet × age interaction F_3,455_ = 1.7, *p* = 0.2; Table S1).

### 3.2. High early-life reproduction is not the cause for decrease in mid-life reproductive output in populations that consume high protein adult diet

To test whether the above mentioned decrease in mid-life reproductive output is due to a tradeoff with investment into early-life reproductive output in HH versus LH i.e. varying developmental diet but high protein adult diet (Table S2; Table S4; Figure 1A), we minimized the investment into reproduction in these populations in early-life by rearing them as virgins throughout early-life, and allowed them to mate and reproduce only in mid-life.

The viable progeny produced in mid-life in these experimental populations were counted to check whether or not difference in mid-life reproductive output between HH and LH exists since their investment into early-life reproduction has been minimized. From Figure 1C, we see that the difference between HH and LH still exists in mid-life despite being reared as virgins in early-life (F_1,57_ = 9.2, *p* = 4×10^−3^; Figure 1C; Table S5).

### 3.3. The effect of developmental and adult diets on ovary size is dependent on age

Effect of developmental diet on ovary size varies with age (developmental diet × age interaction F_3,457_ = 34.5, *p* < 10^−7^; Figure 2A; Table S6). In very-early-life, populations reared on a high protein developmental diet have larger ovaries as compared to populations reared on a low protein developmental diet (for 10th day Tukey’s HSD *p* <10^−7^; Figure 2A; Table S7). However, in mid-life, populations reared in a low protein developmental diet have larger ovaries as compared to populations reared on a high protein developmental diet (for 22nd day Tukey’s HSD *p* = 0.04; Figure 2A; Table S7). In late-life, developmental diet has no effect on ovary size (for 32nd day Tukey’s HSD *p* = 0.89; Figure 2A; Table S7).

**Figure 2.**
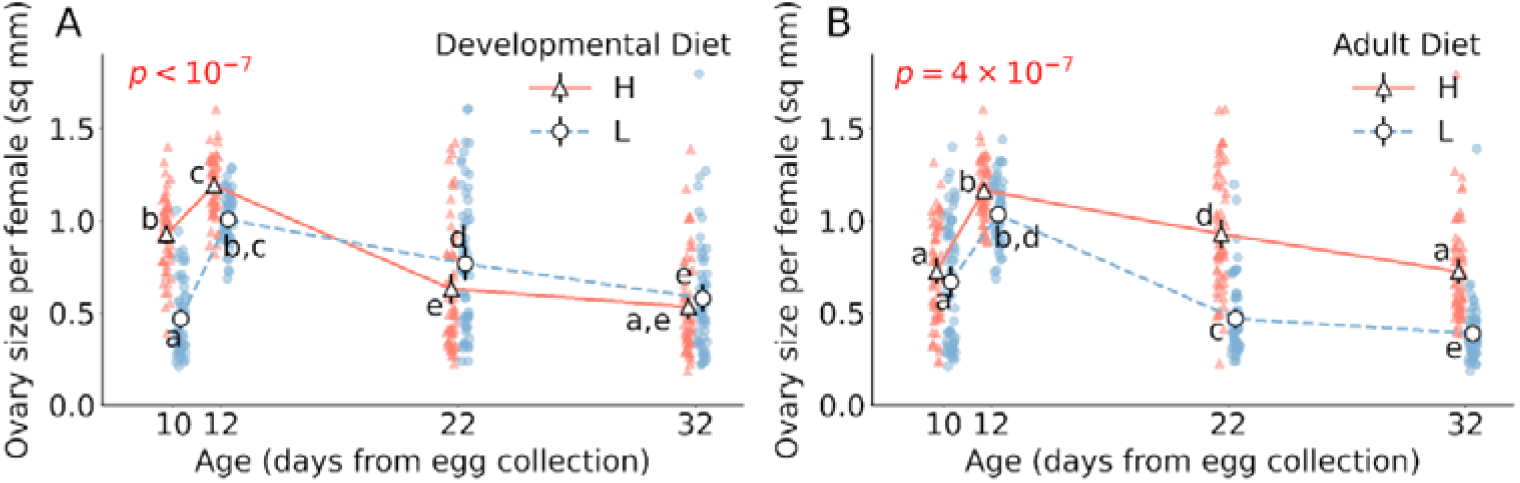
Effect of developmental and adult diet on the size of ovaries in female flies. (A) The effect of developmental diet across age on ovary size. (B) The effect of adult diet across age on ovary size. The triangle marker with solid line represents high protein diet H; the circle marker with dotted line represents low protein diet L. The black solid vertical lines indicate 95% confidence intervalfor each treatment. Some error bars are not visible because of their small size. The p-value in (A) is of developmental diet × age interaction effect. The p-value in (B) is of adult diet × age interaction effect. Different lower-case letters denote statistically significant differences (*p* < 0.05).

Effect of adult diet on ovary size varies with age (adult diet × age interaction F_3,457_ = 9.6, *p* = 3×10^−6^; Figure 2B; Table S6). Populations reared on high protein adult diet have larger ovaries than populations reared in low protein adult diet at all measured age point with statistical significance seen in mid and late-life (for 10th day Tukey’s HSD *p* = 0.90, for 12th day Tukey’s HSD *p* = 0.49, for 22nd day Tukey’s HSD *p* < 10^−7^, for 32nd day Tukey’s HSD *p* < 10^−7^; Figure 2B; Table S8).

We find that the effect of developmental diet on ovary size is independent of adult diet and vice versa (developmental diet × adult diet interaction F_1,457_ = 3.7, *p* = 0.06; Table S6). The three-way interaction of developmental diet, adult diet and age is also not significant (developmental diet × adult diet × age interaction F_3,457_ = 0.6, *p* = 0.6; Table S6).

### 3.4. High protein developmental diet increases the number of ovarioles

Irrespective of adult diet and age, populations reared in high protein developmental diet have a greater number of ovarioles (F_1,472_ = 5.1, *p* = 0.02; Figure 3C; Table S9). The number of ovarioles do not vary due to adult diet (F_1,472_ = 3.5, *p* = 0.06; Table S9). There is also no change in the number of ovarioles with age in our measured age-points (F_3,472_ = 1.7, *p* = 0.16; Table S9).

**Figure 3.**
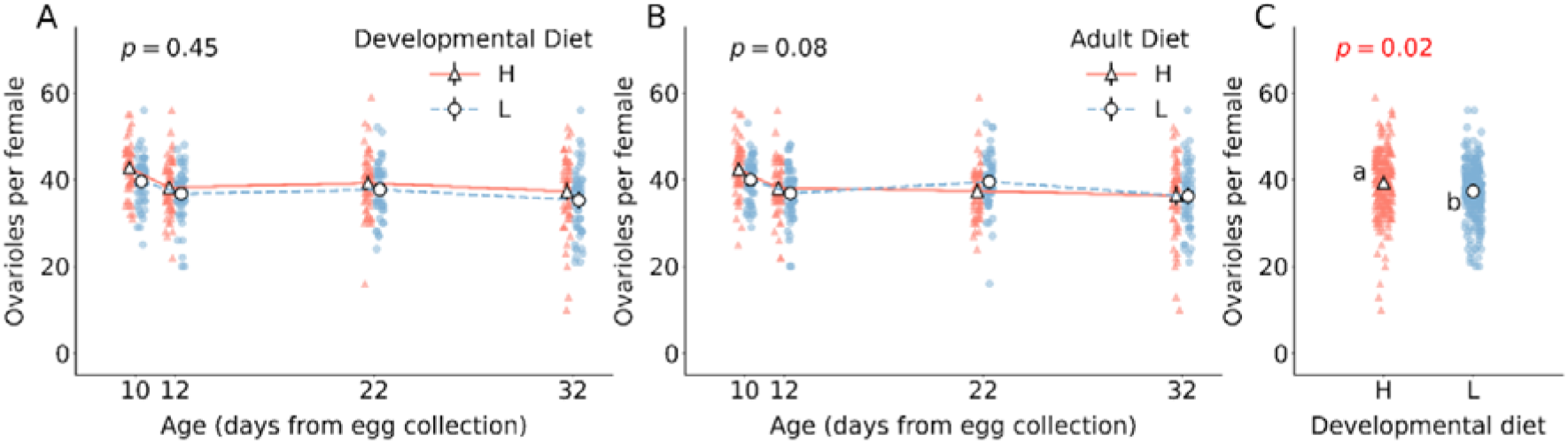
Effect of developmental and adult diet on the total number of ovaries per female flies. (A) The effect of developmental diet across age on the number of ovarioles. (B) The effect of adult diet across age on the number of ovarioles. (C) The effect of developmental diet on the number of ovarioles, irrespective of adult diet and age. The triangle marker with solid line represents high protein diet H; the circle marker with dotted line represents low protein diet L. The black solid vertical lines indicate 95% confidence interval for each treatment. Some error bars are not visible because of their small size. In (A) and (B), the interaction effect is not significant. The p-value in (A) is of developmental diet × age interaction effect. The p-value in (B) is of adult diet × age interaction effect. The p-value in (C) is the main effect of developmental diet. Different lower-case letters in (C) denote statistically significant differences (*p* < 0.05).

### 3.5. Total number of egg chambers in ovaries varies with age, but is independent of diet

The number of egg chambers vary with age (F_3,304_ = 10.7, *p* = 9×10^−7^; Figure 4C; Table S10); after the initial increase in number in early-life, the number of egg chambers decreases with age (Figure 4C; Table S11). However, the number of egg chambers do not vary in response to either developmental diet (F_1,304_ = 3.3, *p* = 0.07; Table S10) or adult diet (F_1,304_ = 1.3, *p* = 0.25; Table S10).

**Figure 4.**
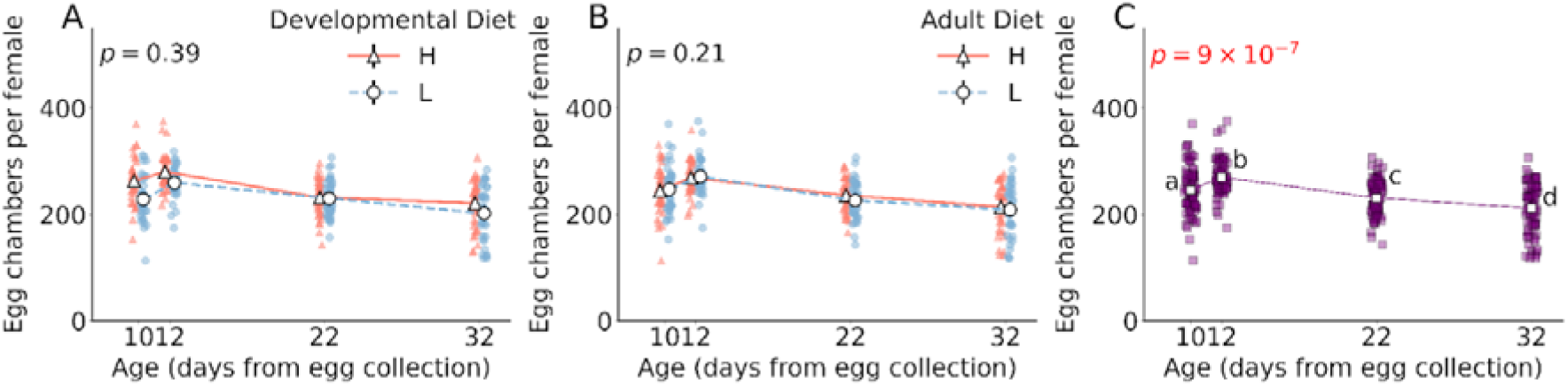
Effect of developmental diet, adult diet and age on the total number of egg chambers per female flies. (A) The effect of developmental diet across age on the number of egg chambers. (B) The effect of adult diet across age on the number of egg chambers. (C) The change in the number of egg chambers with age, irrespective of differences in developmental and adult diets. The triangle marker with solid line represents high protein diet H; the circle marker with dotted line represents low protein diet L. The black solid vertical lines indicate 95% confidence interval for each treatment. Some error bars are not visible because of their small size. In (A) and (B), the interaction effect is not significant. The p-value in (A) is of developmental diet × age interaction effect. The p-value in (B) is of adult diet × age interaction effect. The p-value in (C) is the main effect of age. Different lower-case letters in (C) denote statistically significant differences (*p* < 0.05).

### 3.6. The effect of developmental and adult diets on the proportion of egg chambers that undergo vitellogenesis is dependent on age

Effect of developmental diet on proportion of egg chambers that undergo vitellogenesis varies with age (developmental diet × age interaction F_3,304_ = 9.3, *p* = 7×10^−6^; Figure 5A; Table S12). In very-early-life, populations reared in a high protein developmental diet have a greater proportion of egg chambers that undergo vitellogenesis as compared to populations reared in a low protein developmental diet (for 10th day Tukey’s HSD *p* = 1×10^−5^; Figure 5A; Table S13). However, we find that developmental diet has no effect on the proportion of egg chambers that undergo vitellogenesis at any other age-points (for 12th day Tukey’s HSD *p* = 0.54, for 22nd day Tukey’s HSD *p* = 0.61, for 32nd day Tukey’s HSD *p* = 0.98; Figure 5A; Table S13).

**Figure 5.**
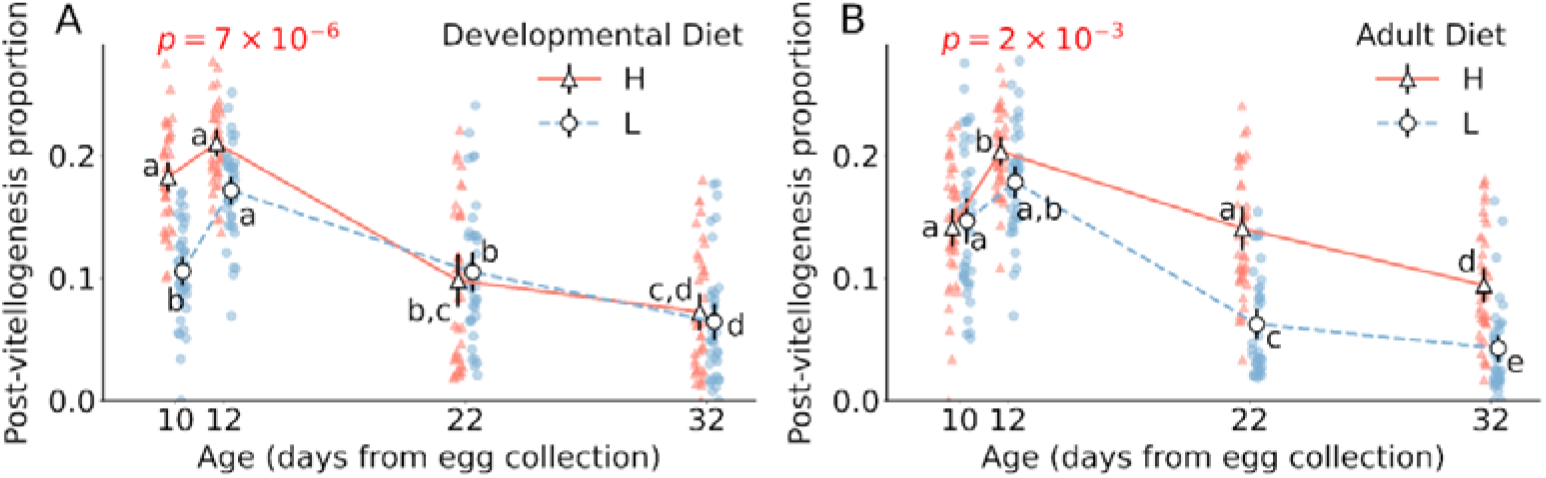
Effect of developmental and adult diet on the proportion of egg chambers that underwent vitellogenesis. (A) The effect of developmental diet across age on the proportion of egg chambers that are in stages >= 8 i.e. egg chambers that undergo vitellogenesis. (B) The effect of adult diet across age on the proportion of egg chambers that are in stages >= 8 i.e. egg chambers that undergo vitellogenesis. The triangle marker with solid line represents high protein diet H; the circle marker with dotted line represents low protein diet L. The black solid vertical lines indicate 95% confidence interval for each treatment. Some error bars are not visible because of their small size. The p-value in (A) is of developmental diet × age interaction effect. The p-value in (B) is of adult diet × age interaction effect. Different lower-case letters denote statistically significant differences (*p* < 0.05).

Effect of adult diet on proportion of egg chambers that undergo vitellogenesis varies with age (adult diet × age interaction F_3,304_ = 5.0, *p* = 2×10^−3^; Figure 5B; Table S12). Populations reared in high protein adult diet have greater proportion of egg chambers that undergo vitellogenesis than populations reared in low protein adult diet at all measured age points, except in very-early and early-life (for 10th day Tukey’s HSD *p* = 1.0, for 12th day Tukey’s HSD *p* = 0.92, for 22nd day Tukey’s HSD *p* < 10^−7^, for 32nd day Tukey’s HSD *p* < 10^−7^; Figure 5B; Table S14).

The effect of developmental diet on the proportion of egg chambers that undergo vitellogenesis is independent of adult diet and vice versa (developmental diet × adult diet interaction F_1,304_ = 0.3, *p* = 0.6; Table S12). The three-way interaction of developmental diet, adult diet and age is also not significant (developmental diet × adult diet × age interaction F_3,304_ = 2.5, *p* = 0.06; Table S12).

## DISCUSSION

Diet significantly influences reproductive outcomes across various species (Lee *et al*., 2008; Millon *et al*., 2011; Wong and Kölliker, 2014; Pigeon *et al*., 2019; Duxbury and Chapman, 2020;). However, the nuanced effects of diet composition at distinct life stages on reproductive output and its aging remain inadequately explored. This has led to conflicting results in the literature regarding the effect of developmental diet on reproductive output (May *et al*., 2015; Duxbury and Chapman, 2020). In this study, we empirically addressed these issues by employing isocaloric diets with differing protein-to-carbohydrate ratios during both the developmental and adult phases in a full factorial manner in a large, outbred population of the common fruit fly, *Drosophila melanogaster*, and subsequently measuring their reproductive output across various ages. We further probed into the sub-organismal levels to explore the underlying mechanisms behind the observed diet-induced modulation of reproductive senescence.

We found that the effect of developmental and adult diets on reproductive output of populations vary with age. Our results reveal that there is a gradual shift in reproductive output being affected only by developmental diet in very-early-life to reproductive output being affected by both development and adult diets in early-life and mid-life to reproductive output being affected only by adult diet in late-life. This progression aligns with results from studies that have looked at the effect of developmental diet across life and showed that the impact of development diet on reproductive output and gene expression is pronounced in early-life, but diminishes significantly in later stages (May *et al*., 2015, 2021). This age-dependent trend potentially clarifies the previously noted inconsistencies in experimental data regarding various hypotheses on the impact of developmental diet on adult phenotypic traits, such as reproductive output (May *et al*., 2015; Duxbury and Chapman, 2020) as elaborated below.

In very-early-life, reproductive output was found to be exclusively influenced by the developmental diet. Our observations revealed that populations reared on a high-protein developmental diet produced a greater number of viable offspring compared to those on a low-protein developmental diet (Figure 1A). In early-life, the reproductive output was shaped by both the developmental and adult diets. Specifically, populations raised on a high-protein regimen during both developmental and adult phases had higher progeny viability than those raised on low-protein diets during these respective stages (Figures 1A and 1B). Consequently, in both very-early and early-life, the impact of the developmental diet on reproductive output aligns with the silver spoon hypothesis, which posits a positive correlation between developmental conditions and subsequent adult-stage performance (Grafen, 1988). In mid-life, although reproductive output was influenced by both developmental and adult diets, interestingly, we observed that populations reared in low protein developmental diet and high protein adult diet produce more viable progeny than populations reared in high protein developmental diet and low protein adult diet, respectively (Figures 1A and 1B).

Consequently, the impact of the developmental diet on reproductive output in mid-life did not conform to the predictions of the silver spoon, thrifty phenotype, or PAR hypotheses. In contrast, during late-life, the reproductive output was solely influenced by the adult diet, with populations fed a high-protein adult diet producing more viable progeny than those on a low-protein adult diet. Our results align with previous research on collared flycatcher females, where ‘silver-spoon’ individuals exhibited enhanced early-life reproductive output at the expense of accelerated reproductive aging and associated fitness declines in later life (Spagopoulou *et al*., 2020). This age-specific trend highlights the intricate relationship between diet composition, reproductive output and aging, potentially explaining the discrepancies observed across studies on the impact of developmental nutrition on fertility.

We further explored how developmental and adult diets impact reproductive output at the suborganismal level, leading to above patterns of reproductive senescence in these flies. We systematically examined the ovaries, ovarioles, and egg chambers under various dietary conditions across different ages. It is well-documented that diet significantly influences ovarian function (Armstrong, 2020), with ovarian size reflecting reproductive output.

Consistent with this, we found that changes in ovary size due to diet closely mirrored the overall age-dependent reproductive output (Figure 2).

In *Drosophila melanogaster*, ovary development occurs during the late larval and early pupal stages, with each ovary containing approximately 18 egg-producing chambers called ovarioles. The formation of these ovarioles is contingent on the presence of terminal filament (TF) cells in the larva (Green and Extavour, 2014), which in turn depends on the number of somatic gonad precursors (SGP cells). SGP cell proliferation is influenced by the Insulin Receptor (InR) pathway, a crucial component of the canonical nutrient-sensing insulin/insulin-like growth factor signalling (IIS) (Green and Extavour, 2014). IIS is upregulated by a high protein diet through elevated expression of DILPs in insulin-producing cells (IPCs) (Lin *et al*., 2018). These insights corroborate our findings that the dietary composition during the larval stage affects the number of ovarioles, specifically with a high-protein developmental diet leading to an increase in the number of ovarioles formed (Figure 3C). Notably, neither the adult diet nor age significantly alters the ovariole count, suggesting that the ovarioles, once formed, persist into later life stages. Therefore, the diet and age-induced variations in reproductive output are likely governed by mechanisms operating at lower levels of organization within the ovarioles.

Within the ovarioles, germ stem cells (GSCs) reside at the anterior tip, dividing asymmetrically to form egg chambers. As the egg chambers leave the germarium and continue to move within the ovariole from the anterior tip to the posterior end, they grow in size, mature and produce fully developed oocytes competent for fertilization. These egg chambers are categorized into 14 distinct stages based on their morphology, with a critical checkpoint occurring between stages 7 to 8, marked by vitellogenesis – the initiation of increased accumulation of yolk by the oocyte (Cummings and King, 1969; King, 1970).

Nutrition plays a significant role in this transition; malnutrition reduces yolk protein transcription, limiting the number of egg chambers capable of undergoing vitellogenesis (Bownes, 1989). Failure to undergo vitellogenesis results in the termination of the egg chambers (Terashima and Bownes, 2006), which ultimately reduces the reproductive output of the individuals. To discern whether dietary regulation of reproductive output operates through the production of egg chambers or via the vitellogenesis checkpoint of oogenesis, we quantified both the total number of egg chambers and the proportion of egg chambers that underwent vitellogenesis. Notably, the total egg chamber count did not vary in response to diet, suggesting a consistent rate of egg chamber production within each ovary across dietary regimes (Figures 4A and 4B). The total egg chamber numbers, however, varied with age, peaking from very-early to early-life before gradually declining in mid- and late-life (Figure 4C). This pattern indicates an initial surge in egg chamber production, followed by an age-related decrease, typically linked to the diminishing pool of germline stem cells (GSCs) and alterations in the ovarian stem cell niche (Miller *et al*., 2014). Moreover, interestingly, the impact of developmental and adult diets on the proportion of egg chambers that underwent vitellogenesis (stage 8 and beyond) mirrors their effects on ovary size and overall reproductive success. In the initial stages of life, this proportion is influenced solely by the developmental diet, whereas in mid-to late-life, the adult diet predominates (Figures 5A and 5B). For both life phases, a high-protein diet significantly increases the proportion of egg chambers that undergo vitellogenesis, corroborating with existing research that demonstrates that *Drosophila* insulin-like peptides (DILPs), which are upregulated in response to dietary amino acids, directly facilitate vitellogenesis (LaFever and Drummond-Barbosa, 2005).

Given the extensive histolysis of larval structures during pupation in holometabolous organisms like *Drosophila*, it is interesting to note that the effect of developmental diet persists into adulthood to influence reproduction. We hypothesize that certain persistent structures, such as the larval fat body, malpighian tubules, specific gut tissues, and a few neurons that do not undergo histolysis (Riddiford, 1993; Lee *et al*., 1999; Klapper, 2000; Nelliot *et al*., 2006; Aguila *et al*., 2007, 2013), could mediate the influence of developmental diets on reproductive output, particularly the larval fat body which is implicated in oogenesis (Sieber and Spradling, 2015) and yolk protein synthesis (Bownes and Reid, 1990). The larval fat body is present for only about a week into the adult stage (Aguila *et al*., 2007, 2013), which may account for our observation that the effect of developmental diet is limited to very-early and early-life. Future experiments should focus on tracking the larval fat body in adults reared on varied diets to elucidate this relationship. Another aspect to consider is the potential influence of developmental timing differences under isocaloric diets with divergent diet compositions (Mudunuri *et al*., 2024), highlighting the need for experiments matching adults by age to nullify the effects of diet induced plasticity in developmental time.

Furthermore, it is important to note that our experiments do not tease apart male versus female effects since the males and females used in our experiments were housed together in their respective dietary regimens. Future experiments could focus on understanding sexual dimorphism in reproductive senescence due to dietary differences. Additionally, when considering a protein-rich adult diet, it’s intriguing why flies reared on a high-protein diet during development (HH regime) exhibit lower mid-life reproductive output compared to those on a low-protein developmental diet (LH regime). A possible explanation is an early-life reproductive trade-off, where HH flies expend more resources on early reproduction, leading to diminished mid-life output. This aligns with studies suggesting that increased early-life reproduction could reduce later reproductive capacity (Nussey *et al*., 2006; McKenna-Ell *et al*., 2023). To test this, we maintained both HH and LH flies as virgins until mid-life, equalizing early reproductive investment. Yet, upon mating in mid-life, LH flies still produced more offspring (Figure 1C), challenging the early-late-life trade-off hypothesis and indicating the need for further investigation to uncover the underlying mechanisms. Lastly, we ponder whether developmental and adult diets influence reproductive output and senescence via separate mechanisms. This is prompted by our observation that the effects of developmental and adult diets on reproductive output were non-interactive. While exploring this lies beyond the scope of the current study, a systematic investigation into the cellular and molecular mechanisms underlying stage-specific dietary impacts could elucidate this query.

In summary, our research reveals the significant influence of developmental and adult diets on reproductive output and its functional senescence. Through detailed experimentation, we have identified that the effect of developmental diet on reproduction is regulated via ovarioles formation and vitellogenesis; while, the effect of adult diet on reproduction is mostly regulated only via vitellogenesis (Table S15). Furthermore, our findings underscore the importance of considering age when evaluating the impact of developmental and adult diets on reproductive output. The insights obtained from this research holds implications for both public health and animal conservation. In a world facing aging populations, fertility challenges, and lifestyle diseases, our insights can guide improved dietary interventions, enhancing reproductive health and longevity. Simultaneously, as climate change and habitat alteration lead to dietary mismatches in wildlife, our findings offer valuable perspectives for conservation strategies, ensuring the reproductive vitality of animal populations. Bridging nutritional science and aging biology, our research aligns human health objectives with ecological sustainability.

## Acknowledgement

The authors also thank Prof. Sutirth Dey from Indian Institute of Science Education and Research (IISER)-Pune for his valuable comments on the experiment design. The authors thank Bio-Imaging Facility, Trivedi School of Biosciences at Ashoka University for the instrumentation support for the experiments. RBG acknowledges the support from Kishore Vaigyanik Protsahan Yojana (KVPY) fellowship. DK, IF, NK, CS and SC acknowledge the funding support from the Research and Development Office (RDO), Ashoka University. MC acknowledges the funding support from the Department of Biotechnology (DBT), India through the Junior Research Fellowship (JRF). ST acknowledges the support of DBT/Wellcome Trust India Alliance Early Career Fellowship (#IA/E/18/1/504347) for research funding and Ashoka University for the infrastructure.

