## Supplementary for "Effect of developmental and adult diet composition on reproductive aging in *Drosophila melanogaster*"

### Table S1. Statistics of viable progeny count analysis

|  | Sum Sq | Df | F values | Pr(>F) |
| --- | --- | --- | --- | --- |
| Age | 184.1222681 | 3 | 55.66035221 | 0 |
| Larva_diet | 18.66780657 | 1 | 16.92989174 | 4.60459E-05 |
| Adult_diet | 5.159319551 | 1 | 4.679002921 | 0.031053548 |
| Age:Larva_diet | 20.18624794 | 3 | 6.102323646 | 0.000446604 |
| Age:Adult_diet | 71.75352864 | 3 | 21.69116597 | 0 |
| Larva_diet:Adult_diet | 0.330722305 | 1 | 0.299933085 | 0.584192919 |
| Age:Larva_diet:Adult_diet | 5.526270316 | 3 | 1.670597236 | 0.172541313 |
| Residuals | 501.7074013 | 455 |  |  |

*Red highlight indicates statistical significance i.e p < 0.05*

### Table S2. Statistics of viable progeny count Tukey analysis for developmental diet x age

|  | contrast | estimate | standard error | z.ratio | p.value |
| --- | --- | --- | --- | --- | --- |
| 1 | Age10 L - Age12 L | -0.443863801 | 0.083735367 | -5.30079242 | 3.19802E-06 |
| 2 | Age10 L - Age22 L | 0.200187181 | 0.089182075 | 2.244701984 | 0.324882155 |
| 3 | Age10 L - Age32 L | 0.675117247 | 0.092877947 | 7.268864884 | 0 |
| 4 | Age10 L - Age10 H | -0.472813573 | 0.083649719 | -5.652303179 | 4.41165E-07 |
| 5 | Age10 L - Age12 H | -0.759396465 | 0.082869338 | -9.163781042 | 0 |
| 6 | Age10 L - Age22 H | 0.526864366 | 0.09071943 | 5.807624332 | 1.76808E-07 |
| 7 | Age10 L - Age32 H | 0.855884051 | 0.095663447 | 8.946824272 | 0 |
| 8 | Age12 L - Age22 L | 0.644050982 | 0.086977208 | 7.40482476 | 0 |
| 9 | Age12 L - Age32 L | 1.118981048 | 0.090762907 | 12.32861629 | 0 |
| 10 | Age12 L - Age10 H | -0.028949772 | 0.081294943 | -0.356107907 | 0.999965912 |
| 11 | Age12 L - Age12 H | -0.315532664 | 0.080491734 | -3.92006293 | 0.002254234 |
| 12 | Age12 L - Age22 H | 0.970728167 | 0.088552849 | 10.96213366 | 0 |
| 13 | Age12 L - Age32 H | 1.299747852 | 0.093611363 | 13.88450941 | 0 |
| 14 | Age22 L - Age32 L | 0.474930066 | 0.095810938 | 4.956950364 | 1.96968E-05 |
| 15 | Age22 L - Age10 H | -0.673000753 | 0.086894755 | -7.745010102 | 0 |
| 16 | Age22 L - Age12 H | -0.959583646 | 0.086143775 | -11.13932657 | 0 |
| 17 | Age22 L - Age22 H | 0.326677186 | 0.093719997 | 3.485672183 | 0.011548899 |
| 18 | Age22 L - Age32 H | 0.65569687 | 0.098513542 | 6.655905926 | 7.87992E-10 |
| 19 | Age32 L - Age10 H | -1.14793082 | 0.090683896 | -12.65859622 | 0 |
| 20 | Age32 L - Age12 H | -1.434513712 | 0.089964552 | -15.94532163 | 0 |
| 21 | Age32 L - Age22 H | -0.14825288 | 0.097243552 | -1.524552299 | 0.794386886 |
| 22 | Age32 L - Age32 H | 0.180766804 | 0.101871432 | 1.774460233 | 0.637547716 |
| 23 | Age10 H - Age12 H | -0.286582892 | 0.080402631 | -3.564347203 | 0.008732977 |
| 24 | Age10 H - Age22 H | 0.999677939 | 0.088471865 | 11.29938817 | 0 |
| 25 | Age10 H - Age32 H | 1.328697624 | 0.093534758 | 14.20538895 | 0 |
| 26 | Age12 H - Age22 H | 1.286260831 | 0.087734386 | 14.66085185 | 0 |
| 27 | Age12 H - Age32 H | 1.615280516 | 0.092837507 | 17.39900793 | 0 |
| 28 | Age22 H - Age32 H | 0.329019684 | 0.099907409 | 3.293246082 | 0.022162905 |

*Red highlight indicates statistical significance i.e p < 0.05*

### Table S3. Statistics of viable progeny count Tukey analysis for adult diet x age

|  | contrast | estimate | standard error | z.ratio | p.value |
| --- | --- | --- | --- | --- | --- |
| 1 | Age10 L - Age12 L | -0.355006402 | 0.083132234 | -4.270382053 | 0.00051665 |
| 2 | Age10 L - Age22 L | 1.133414028 | 0.094756244 | 11.96136507 | 0 |
| 3 | Age10 L - Age32 L | 1.384181651 | 0.100087776 | 13.82967733 | 0 |
| 4 | Age10 L - Age10 H | -0.228424913 | 0.083649719 | -2.730731383 | 0.113381314 |
| 5 | Age10 L - Age12 H | -0.603865204 | 0.082472419 | -7.322026089 | 0 |
| 6 | Age10 L - Age22 H | -0.161973822 | 0.083895417 | -1.930663523 | 0.529561226 |
| 7 | Age10 L - Age32 H | 0.391208306 | 0.087143414 | 4.48924696 | 0.000192386 |
| 8 | Age12 L - Age22 L | 1.48842043 | 0.09303744 | 15.99808028 | 0 |
| 9 | Age12 L - Age32 L | 1.739188053 | 0.098462086 | 17.66353045 | 0 |
| 10 | Age12 L - Age10 H | 0.126581489 | 0.081697582 | 1.549390889 | 0.780374472 |
| 11 | Age12 L - Age12 H | -0.248858802 | 0.080491734 | -3.091731147 | 0.041731159 |
| 12 | Age12 L - Age22 H | 0.19303258 | 0.081949133 | 2.355517053 | 0.263620844 |
| 13 | Age12 L - Age32 H | 0.746214708 | 0.085271296 | 8.751065595 | 0 |
| 14 | Age22 L - Age32 L | 0.250767623 | 0.108455335 | 2.312174148 | 0.286711499 |
| 15 | Age22 L - Age10 H | -1.361838941 | 0.093500119 | -14.5651038 | 0 |
| 16 | Age22 L - Age12 H | -1.737279233 | 0.092448346 | -18.79189088 | 0 |
| 17 | Age22 L - Age22 H | -1.29538785 | 0.093719997 | -13.82189387 | 0 |
| 18 | Age22 L - Age32 H | -0.742205722 | 0.096638354 | -7.68023972 | 0 |
| 19 | Age32 L - Age10 H | -1.612606564 | 0.098899391 | -16.30552579 | 0 |
| 20 | Age32 L - Age12 H | -1.988046855 | 0.097905638 | -20.30574443 | 0 |
| 21 | Age32 L - Age22 H | -1.546155473 | 0.09910729 | -15.60082482 | 0 |
| 22 | Age32 L - Age32 H | -0.992973345 | 0.101871432 | -9.747319045 | 0 |
| 23 | Age10 H - Age12 H | -0.375440291 | 0.081026085 | -4.633573161 | 9.75528E-05 |
| 24 | Age10 H - Age22 H | 0.066451091 | 0.082474041 | 0.805721294 | 0.992861404 |
| 25 | Age10 H - Age32 H | 0.619633219 | 0.085775877 | 7.223863404 | 0 |
| 26 | Age12 H - Age22 H | 0.441891383 | 0.081279714 | 5.436674918 | 1.5089E-06 |
| 27 | Age12 H - Age32 H | 0.995073511 | 0.08462816 | 11.75818437 | 0 |
| 28 | Age22 H - Age32 H | 0.553182128 | 0.086015502 | 6.431191057 | 3.54186E-09 |

*Red highlight indicates statistical significance i.e p < 0.05*

Table S4. Effect sizes of the number of viable progeny produced compared between the four dietary treatments (HH, HL, LH and LL) at each measured age (10, 12, 22 and 32)

| Age | Groups  (regime 1 - regime 2) | Mean difference  (regime 1 - regime 2) | Effect size  (Cohen’s d) | Interpretation |
| --- | --- | --- | --- | --- |
| 10 | HH-LH | 11.5 | 1.340800264 | Large |
| 10 | HH-HL | 5.483333333 | 0.643431035 | Medium |
| 10 | HH-LL | 16.75057471 | 2.019201605 | Large |
| 10 | LH-HL | -6.016666667 | 0.776498273 | Medium |
| 10 | LH-LL | 5.250574713 | 0.701570741 | Medium |
| 10 | HL-LL | 5.250574713 | 1.518519955 | Large |
| 12 | HH-LH | 19.15 | 2.159882463 | Large |
| 12 | HH-HL | 17.03333333 | 2.250573132 | Large |
| 12 | HH-LL | 21.5 | 2.896495465 | Large |
| 12 | LH-HL | -2.116666667 | 0.289920494 | Small |
| 12 | LH-LL | 2.35 | 0.328684077 | Small |
| 12 | HL-LL | 4.466666667 | 0.818368135 | Large |
| 22 | HH-LH | -9.85 | 0.726714491 | Medium |
| 22 | HH-HL | 14.66666667 | 1.593205791 | Large |
| 22 | HH-LL | 12.84285714 | 1.364598283 | Large |
| 22 | LH-HL | 24.51666667 | 2.261217869 | Large |
| 22 | LH-LL | 22.69285714 | 2.049853936 | Large |
| 22 | HL-LL | -1.823809524 | 0.407290118 | Small |
| 32 | HH-LH | -3.556896552 | 0.371849854 | Small |
| 32 | HH-HL | 7.737547893 | 1.029738235 | Large |
| 32 | HH-LL | 7.11453202 | 0.955326278 | Large |
| 32 | LH-HL | 11.29444444 | 1.38896081 | Large |
| 32 | LH-LL | 10.67142857 | 1.323958452 | Large |
| 32 | HL-LL | -0.623015873 | 0.118226957 | non-significant |

### Table S5. Statistics of viable progeny produced in mid-life by flies reared as virgins in early-life and allowed to mate and reproduce only in mid-life

|  | Sum Sq | Df | F values | Pr(>F) |
| --- | --- | --- | --- | --- |
| Treatment | 6.57109247 | 1 | 9.24966952 | 0.00355596 |
| Residuals | 40.4935841 | 57 |  |  |

*Red highlight indicates statistical significance i.e p < 0.05*

### Table S6. Statistics of ovary size analysis

|  | Sum Sq | Df | F values | Pr(>F) |
| --- | --- | --- | --- | --- |
| Age | 15.81344618 | 3 | 49.24674166 | 0 |
| Larva_diet | 10.80655468 | 1 | 100.9623583 | 0 |
| Adult_diet | 1.387885795 | 1 | 12.96659547 | 0.000351827 |
| Age:Larva_diet | 11.0713314 | 3 | 34.47869559 | 0 |
| Age:Adult_diet | 3.08851079 | 3 | 9.618339433 | 3.61259E-06 |
| Larva_diet:Adult_diet | 0.393838012 | 1 | 3.679508932 | 0.055707863 |
| Age:Larva_diet:Adult_diet | 0.206260416 | 3 | 0.642342807 | 0.588101666 |
| Residuals | 48.91521525 | 457 |  |  |

*Red highlight indicates statistical significance i.e p < 0.05*

### Table S7. Statistics of ovary size Tukey analysis for developmental diet x age

|  | contrast | estimate | standard error | t.ratio | p.value |
| --- | --- | --- | --- | --- | --- |
| 1 | Age10 L - Age12 L | -0.535901355 | 0.048194261 | -11.11960925 | 0 |
| 2 | Age10 L - Age22 L | -0.298442593 | 0.039373238 | -7.579833539 | 0 |
| 3 | Age10 L - Age32 L | -0.111477778 | 0.032562916 | -3.423458031 | 0.015406364 |
| 4 | Age10 L - Age10 H | -0.457814815 | 0.043879012 | -10.43357171 | 0 |
| 5 | Age10 L - Age12 H | -0.724552235 | 0.055537533 | -13.0461725 | 0 |
| 6 | Age10 L - Age22 H | -0.162144445 | 0.034462116 | -4.705005527 | 9.11329E-05 |
| 7 | Age10 L - Age32 H | -0.065131482 | 0.030685204 | -2.122569618 | 0.401885001 |
| 8 | Age12 L - Age22 L | 0.237458763 | 0.055462915 | 4.281397087 | 0.000594551 |
| 9 | Age12 L - Age32 L | 0.424423578 | 0.050854956 | 8.345766264 | 0 |
| 10 | Age12 L - Age10 H | 0.07808654 | 0.058747346 | 1.32919264 | 0.887420828 |
| 11 | Age12 L - Age12 H | -0.18865088 | 0.067899194 | -2.778396435 | 0.103019142 |
| 12 | Age12 L - Age22 H | 0.373756911 | 0.052091462 | 7.175012828 | 0 |
| 13 | Age12 L - Age32 H | 0.470769874 | 0.049673582 | 9.477268441 | 0 |
| 14 | Age22 L - Age32 L | 0.186964815 | 0.042588633 | 4.390016851 | 0.000373485 |
| 15 | Age22 L - Age10 H | -0.159372222 | 0.051757277 | -3.079223448 | 0.045275139 |
| 16 | Age22 L - Age12 H | -0.426109642 | 0.06195051 | -6.878226565 | 5.55871E-10 |
| 17 | Age22 L - Age22 H | 0.136298148 | 0.044057752 | 3.09362466 | 0.043402572 |
| 18 | Age22 L - Age32 H | 0.233311111 | 0.04117074 | 5.666915646 | 7.16498E-07 |
| 19 | Age32 L - Age10 H | -0.346337037 | 0.046785761 | -7.402616367 | 0 |
| 20 | Age32 L - Age12 H | -0.613074457 | 0.057861536 | -10.5955442 | 0 |
| 21 | Age32 L - Age22 H | -0.050666667 | 0.038094319 | -1.330032088 | 0.887086402 |
| 22 | Age32 L - Age32 H | 0.046346296 | 0.034714859 | 1.335056428 | 0.885072199 |
| 23 | Age10 H - Age12 H | -0.26673742 | 0.064907484 | -4.109501789 | 0.001212357 |
| 24 | Age10 H - Age22 H | 0.29567037 | 0.048126929 | 6.143553671 | 4.90389E-08 |
| 25 | Age10 H - Age32 H | 0.392683333 | 0.045498853 | 8.630620538 | 0 |
| 26 | Age12 H - Age22 H | 0.562407791 | 0.058951261 | 9.540216449 | 0 |
| 27 | Age12 H - Age32 H | 0.659420754 | 0.056826011 | 11.60420631 | 0 |
| 28 | Age22 H - Age32 H | 0.097012963 | 0.036502265 | 2.657724468 | 0.138549809 |

*Red highlight indicates statistical significance i.e p < 0.05*

### Table S8. Statistics of ovary size Tukey analysis for adult diet x age

|  | contrast | estimate | standard error | t.ratio | p.value |
| --- | --- | --- | --- | --- | --- |
| 1 | Age10 L - Age12 L | -0.36214133 | 0.054382162 | -6.659193359 | 2.21108E-09 |
| 2 | Age10 L - Age22 L | 0.199485185 | 0.036366396 | 5.485426273 | 1.89686E-06 |
| 3 | Age10 L - Age32 L | 0.279646296 | 0.034567372 | 8.08989162 | 0 |
| 4 | Age10 L - Age10 H | -0.057007407 | 0.043879012 | -1.299195339 | 0.898976183 |
| 5 | Age10 L - Age12 H | -0.497504853 | 0.059207314 | -8.402759996 | 0 |
| 6 | Age10 L - Age22 H | -0.259264815 | 0.049711123 | -5.215428672 | 7.67917E-06 |
| 7 | Age10 L - Age32 H | -0.055448148 | 0.043160667 | -1.284691629 | 0.904287348 |
| 8 | Age12 L - Age22 L | 0.561626515 | 0.049267731 | 11.39948007 | 0 |
| 9 | Age12 L - Age32 L | 0.641787626 | 0.047955164 | 13.38307644 | 0 |
| 10 | Age12 L - Age10 H | 0.305133922 | 0.055046909 | 5.543161786 | 1.39568E-06 |
| 11 | Age12 L - Age12 H | -0.135363524 | 0.067899194 | -1.993595427 | 0.487323684 |
| 12 | Age12 L - Age22 H | 0.102876515 | 0.059799584 | 1.720355025 | 0.673963429 |
| 13 | Age12 L - Age32 H | 0.306693182 | 0.054476029 | 5.629874038 | 8.75896E-07 |
| 14 | Age22 L - Age32 L | 0.080161111 | 0.025783581 | 3.108998421 | 0.041477215 |
| 15 | Age22 L - Age10 H | -0.256492593 | 0.037353145 | -6.866693325 | 5.98445E-10 |
| 16 | Age22 L - Age12 H | -0.696990038 | 0.054547189 | -12.7777443 | 0 |
| 17 | Age22 L - Age22 H | -0.45875 | 0.044057752 | -10.41246952 | 0 |
| 18 | Age22 L - Age32 H | -0.254933333 | 0.036506616 | -6.983209125 | 2.82037E-10 |
| 19 | Age32 L - Age10 H | -0.336653704 | 0.035604015 | -9.455498193 | 0 |
| 20 | Age32 L - Age12 H | -0.77715115 | 0.053364636 | -14.56303675 | 0 |
| 21 | Age32 L - Age22 H | -0.538911111 | 0.042584903 | -12.65498032 | 0 |
| 22 | Age32 L - Age32 H | -0.335094445 | 0.034714859 | -9.652766847 | 0 |
| 23 | Age10 H - Age12 H | -0.440497446 | 0.059818465 | -7.363904192 | 0 |
| 24 | Age10 H - Age22 H | -0.202257407 | 0.05043747 | -4.010062487 | 0.001806598 |
| 25 | Age10 H - Age32 H | 0.001559259 | 0.043995294 | 0.035441501 | 1 |
| 26 | Age12 H - Age22 H | 0.238240038 | 0.064218976 | 3.709807517 | 0.005666309 |
| 27 | Age12 H - Age32 H | 0.442056705 | 0.059293544 | 7.455393589 | 0 |
| 28 | Age22 H - Age32 H | 0.203816667 | 0.049813793 | 4.09157093 | 0.001303719 |

*Red highlight indicates statistical significance i.e p < 0.05*

### Table S9. Statistics of ovariole count analysis

|  | Sum Sq | Df | F values | Pr(>F) |
| --- | --- | --- | --- | --- |
| Age | 5.426794108 | 3 | 1.734958397 | 0.158954631 |
| Larva_diet | 5.366664255 | 1 | 5.147204239 | 0.023734553 |
| Adult_diet | 3.679419674 | 1 | 3.52895647 | 0.060921841 |
| Age:Larva_diet | 2.782942898 | 3 | 0.889713163 | 0.446304793 |
| Age:Adult_diet | 7.227416149 | 3 | 2.310621351 | 0.075530165 |
| Larva_diet:Adult_diet | 0.417764673 | 1 | 0.400680943 | 0.527044353 |
| Age:Larva_diet:Adult_diet | 1.886405902 | 3 | 0.60308825 | 0.61325716 |
| Residuals | 492.1245419 | 472 |  |  |

*Red highlight indicates statistical significance i.e p < 0.05*

### Table S10. Statistics of total number of egg chambers analysis

|  | Sum Sq | Df | F values | Pr(>F) |
| --- | --- | --- | --- | --- |
| Age | 32.7745394 | 3 | 10.7480422 | 9.85E-07 |
| Larva_diet | 3.35437909 | 1 | 3.30009288 | 0.070260667 |
| Adult_diet | 1.32614443 | 1 | 1.30468253 | 0.254258509 |
| Age:Larva_diet | 3.05159834 | 3 | 1.00073741 | 0.39280036 |
| Age:Adult_diet | 4.57140451 | 3 | 1.49914078 | 0.214819233 |
| Larva_diet:Adult_diet | 1.5508089 | 1 | 1.5257111 | 0.217711034 |
| Age:Larva_diet:Adult_diet | 4.11664552 | 3 | 1.35000768 | 0.258255327 |
| Residuals | 309.000771 | 304 |  |  |

*Red highlight indicates statistical significance i.e p < 0.05*

### Table S11. Statistics of total number of egg chambers Tukey analysis of age

|  | contrast | estimate | SE | z.ratio | p.value |
| --- | --- | --- | --- | --- | --- |
| 1 | Age10 - Age12 | -0.0955729 | 0.02589544 | -3.6907232 | 0.001279 |
| 2 | Age10 - Age22 | 0.0593743 | 0.02604457 | 2.27971918 | 0.10274425 |
| 3 | Age10 - Age32 | 0.14889751 | 0.02614321 | 5.6954568 | 7.37E-08 |
| 4 | Age12 - Age22 | 0.15494721 | 0.02595313 | 5.97027086 | 1.42E-08 |
| 5 | Age12 - Age32 | 0.24447042 | 0.02605212 | 9.38389893 | 3.81E-14 |
| 6 | Age22 - Age32 | 0.08952321 | 0.02620035 | 3.41687097 | 0.00354336 |

*Red highlight indicates statistical significance i.e p < 0.05*

### Table S12. Statistics of proportion of egg chambers that undergo vitellogenesis analysis

|  | Sum Sq | Df | F values | Pr(>F) |
| --- | --- | --- | --- | --- |
| Age | 18.0578967 | 3 | 26.7634219 | 2.18E-15 |
| Larva_diet | 3.58789726 | 1 | 15.9527562 | 8.15E-05 |
| Adult_diet | 0.01112342 | 1 | 0.0494577 | 0.82415895 |
| Age:Larva_diet | 6.27983446 | 3 | 9.30727767 | 6.62E-06 |
| Age:Adult_diet | 3.36694766 | 3 | 4.99011829 | 0.00215433 |
| Larva_diet:Adult_diet | 0.06114059 | 1 | 0.27184751 | 0.60247475 |
| Age:Larva_diet:Adult_diet | 1.66189051 | 3 | 2.46307073 | 0.06256847 |
| Residuals | 68.3719322 | 304 |  |  |

*Red highlight indicates statistical significance i.e p < 0.05*

### Table S13. Statistics of proportion of egg chambers that undergo vitellogenesis Tukey analysis for developmental diet x age

|  | contrast | estimate | SE | t.ratio | p.value |
| --- | --- | --- | --- | --- | --- |
| 1 | Age10 L - Age12 L | -0.4837534 | 0.10604425 | -4.561807 | 0.00019678 |
| 2 | Age10 L - Age22 L | 0.03944173 | 0.10604425 | 0.37193649 | 0.99995294 |
| 3 | Age10 L - Age32 L | 0.56275149 | 0.10604425 | 5.30676089 | 5.94E-06 |
| 4 | Age10 L - Age10 H | -0.5481839 | 0.10604425 | -5.1693878 | 1.17E-05 |
| 5 | Age10 L - Age12 H | -0.6878537 | 0.10604425 | -6.4864776 | 9.94E-09 |
| 6 | Age10 L - Age22 H | 0.23236086 | 0.10604425 | 2.19116879 | 0.36000048 |
| 7 | Age10 L - Age32 H | 0.45862326 | 0.10604425 | 4.32482896 | 0.00054355 |
| 8 | Age12 L - Age22 L | 0.52319514 | 0.10604425 | 4.93374345 | 3.63E-05 |
| 9 | Age12 L - Age32 L | 1.04650491 | 0.10604425 | 9.86856785 | 1.11E-12 |
| 10 | Age12 L - Age10 H | -0.0644305 | 0.10604425 | -0.6075808 | 0.99876642 |
| 11 | Age12 L - Age12 H | -0.2041003 | 0.10604425 | -1.9246706 | 0.53500833 |
| 12 | Age12 L - Age22 H | 0.71611427 | 0.10604425 | 6.75297575 | 2.06E-09 |
| 13 | Age12 L - Age32 H | 0.94237667 | 0.10604425 | 8.88663592 | 1.11E-12 |
| 14 | Age22 L - Age32 L | 0.52330977 | 0.10604425 | 4.9348244 | 3.61E-05 |
| 15 | Age22 L - Age10 H | -0.5876256 | 0.10604425 | -5.5413242 | 1.80E-06 |
| 16 | Age22 L - Age12 H | -0.7272954 | 0.10604425 | -6.8584141 | 1.09E-09 |
| 17 | Age22 L - Age22 H | 0.19291913 | 0.10604425 | 1.81923231 | 0.60746972 |
| 18 | Age22 L - Age32 H | 0.41918153 | 0.10604425 | 3.95289248 | 0.00241583 |
| 19 | Age32 L - Age10 H | -1.1109354 | 0.10604425 | -10.476149 | 1.11E-12 |
| 20 | Age32 L - Age12 H | -1.2506052 | 0.10604425 | -11.793238 | 1.04E-12 |
| 21 | Age32 L - Age22 H | -0.3303906 | 0.10604425 | -3.1155921 | 0.0416121 |
| 22 | Age32 L - Age32 H | -0.1041282 | 0.10604425 | -0.9819319 | 0.97672288 |
| 23 | Age10 H - Age12 H | -0.1396698 | 0.10604425 | -1.3170899 | 0.89190859 |
| 24 | Age10 H - Age22 H | 0.78054472 | 0.10604425 | 7.36055655 | 4.93E-11 |
| 25 | Age10 H - Age32 H | 1.00680712 | 0.10604425 | 9.49421672 | 1.11E-12 |
| 26 | Age12 H - Age22 H | 0.92021453 | 0.10604425 | 8.6776464 | 1.12E-12 |
| 27 | Age12 H - Age32 H | 1.14647693 | 0.10604425 | 10.8113066 | 1.09E-12 |
| 28 | Age22 H - Age32 H | 0.2262624 | 0.10604425 | 2.13366017 | 0.39570379 |

*Red highlight indicates statistical significance i.e p < 0.05*

### Table S14. Statistics of proportion of egg chambers that undergo vitellogenesis Tukey analysis for adult diet x age

|  | contrast | estimate | SE | t.ratio | p.value |
| --- | --- | --- | --- | --- | --- |
| 1 | Age10 L - Age12 L | -0.234723 | 0.10604425 | -2.2134442 | 0.34659296 |
| 2 | Age10 L - Age22 L | 0.84192362 | 0.10604425 | 7.93936113 | 2.22E-12 |
| 3 | Age10 L - Age32 L | 1.19093221 | 0.10604425 | 11.230521 | 1.07E-12 |
| 4 | Age10 L - Age10 H | 0.02194137 | 0.10604425 | 0.20690771 | 0.99999917 |
| 5 | Age10 L - Age12 H | -0.3667588 | 0.10604425 | -3.4585449 | 0.01421278 |
| 6 | Age10 L - Age22 H | 4.20E-06 | 0.10604425 | 3.96E-05 | 1 |
| 7 | Age10 L - Age32 H | 0.40056777 | 0.10604425 | 3.77736427 | 0.00466414 |
| 8 | Age12 L - Age22 L | 1.07664666 | 0.10604425 | 10.1528053 | 1.11E-12 |
| 9 | Age12 L - Age32 L | 1.42565525 | 0.10604425 | 13.4439652 | 1.04E-12 |
| 10 | Age12 L - Age10 H | 0.25666441 | 0.10604425 | 2.4203519 | 0.23527282 |
| 11 | Age12 L - Age12 H | -0.1320358 | 0.10604425 | -1.2451007 | 0.91762645 |
| 12 | Age12 L - Age22 H | 0.23472724 | 0.10604425 | 2.2134838 | 0.34656934 |
| 13 | Age12 L - Age32 H | 0.63529081 | 0.10604425 | 5.99080847 | 1.64E-07 |
| 14 | Age22 L - Age32 L | 0.34900859 | 0.10604425 | 3.29115991 | 0.02441546 |
| 15 | Age22 L - Age10 H | -0.8199822 | 0.10604425 | -7.7324534 | 5.47E-12 |
| 16 | Age22 L - Age12 H | -1.2086824 | 0.10604425 | -11.397906 | 1.06E-12 |
| 17 | Age22 L - Age22 H | -0.8419194 | 0.10604425 | -7.9393215 | 2.22E-12 |
| 18 | Age22 L - Age32 H | -0.4413558 | 0.10604425 | -4.1619969 | 0.00106111 |
| 19 | Age32 L - Age10 H | -1.1689908 | 0.10604425 | -11.023613 | 1.08E-12 |
| 20 | Age32 L - Age12 H | -1.557691 | 0.10604425 | -14.689066 | 1.04E-12 |
| 21 | Age32 L - Age22 H | -1.190928 | 0.10604425 | -11.230481 | 1.07E-12 |
| 22 | Age32 L - Age32 H | -0.7903644 | 0.10604425 | -7.4531568 | 2.78E-11 |
| 23 | Age10 H - Age12 H | -0.3887002 | 0.10604425 | -3.6654526 | 0.00698063 |
| 24 | Age10 H - Age22 H | -0.0219372 | 0.10604425 | -0.2068681 | 0.99999917 |
| 25 | Age10 H - Age32 H | 0.3786264 | 0.10604425 | 3.57045656 | 0.00973018 |
| 26 | Age12 H - Age22 H | 0.36676301 | 0.10604425 | 3.45858452 | 0.01421091 |
| 27 | Age12 H - Age32 H | 0.76732658 | 0.10604425 | 7.23590918 | 1.07E-10 |
| 28 | Age22 H - Age32 H | 0.40056357 | 0.10604425 | 3.77732466 | 0.00466481 |

*Red highlight indicates statistical significance i.e p < 0.05*

### Table S15. Summary table of results

| Age | Viable Progeny Count | | Ovary Size | | Number of Ovarioles | | Number of Egg chamber | | Proportion vitellogenesis | |
| --- | --- | --- | --- | --- | --- | --- | --- | --- | --- | --- |
|  | Dev. diet | Adult diet | Dev. diet | Adult diet | Dev. diet | Adult diet | Dev. diet | Adult diet | Dev. diet | Adult diet |
| Very-early-life | H > L | H ~ L | H > L | H ~ L | H > L | H ~ L | H ~ L | | H > L | H ~ L |
| Early-life | H > L | H > L | H ~ L | H ~ L |  |  |  |  | H ~ L | H ~ L |
| Mid-life | H < L | H > L | H < L | H > L |  |  |  |  | H ~ L | H > L |
| Late-life | H ~ L | H > L | H ~ L | H > L |  |  |  |  | H ~ L | H > L |

*NOTE: H diet is a protein-to-carbohydrate ratio of 0.7; and, L diet is a protein-to-carbohydrate ratio of 0.25*
